# There is no convincing evidence that *Methylobacterium extorquens* AM1 can produce N-deoxyschizokinen A

**DOI:** 10.64898/2026.07.03.736418

**Authors:** Sophie M. Gutenthaler-Tietze, Patrick Weis, Lena J. Daumann

**Affiliations:** Chair of Bioinorganic Chemistry, Heinrich Heine University of Düsseldorf, Universitätsstr. 1, 40225 Düsseldorf, Germany; Institut für Physikalische Chemie, Karlsruher Institut für Technologie, Fritz-Haber-Weg 2, 76131 Karlsruhe, Germany

**Keywords:** Metallophores, siderophores, lanthanophore, lanthanides, iron, methylotrophy, C1 metabolism, metal-uptake, bacterial growth

## Abstract

It was recently reported that *Methylobacterium extorquens* AM1 produces the citrate– hydroxamate siderophore *N*-deoxyschizokinen A, identified by LC-HRMS. Multiple properties were inconsistent with the assignment: the feature eluted far later than the other schizokinen derivatives (17 min versus 6-8 min), a reversed-phase shift larger than a single-hydroxyl difference in a molecule can explain, further its accurate mass deviated from the calculated one by 28 ppm, well outside the error on the co-analyzed standards and its diagnostic *m/z* 105 and 77 fragments suggest a molecule with an aromatic moiety. A replicate comparison of identical samples in plastic versus glass autosampler vials was decisive: the *m/z* 387 feature was reproducibly present with plastic vials and absent with glass. We therefore conclude that the reported detection of *N*-deoxyschizokinen A in *M. extorquens* AM1 is an artifact, and recommend glass-vial and solvent-blank controls, an explicit accurate-mass threshold, and narrow MS/MS isolation when assigning trace siderophore-like features from complex extracts.

## Introduction

How poorly bioavailable Ln are taken up by bacteria is an active area of research.^[1–6]^ Early on, the involvement of small chelators (lanthanophores) similar to siderophores in Ln-uptake was suggested and recently, the first chelator involved in Ln-dependent growth has been isolated and structurally characterized from the methylotroph *Methylobacterium extorquens* AM1 and also PA1.^[1–4,7]^ In a recent preprint on Biorxiv,^[8]^ it was stated that a specific chelator was found in the extract of spent *M. extorquens* AM1 medium and assigned to the siderophore N-deoxyschizokinen A (NDSA). There are several inconsistencies with that assignment, which was done solely with LC-MS. Since we are actively working on this topic it was important to us to set the record straight and thus we gathered evidence to show that AM1 does not produce the siderophore N-deoxyschizokinen A.^[8,9]^

**Figure 1.**
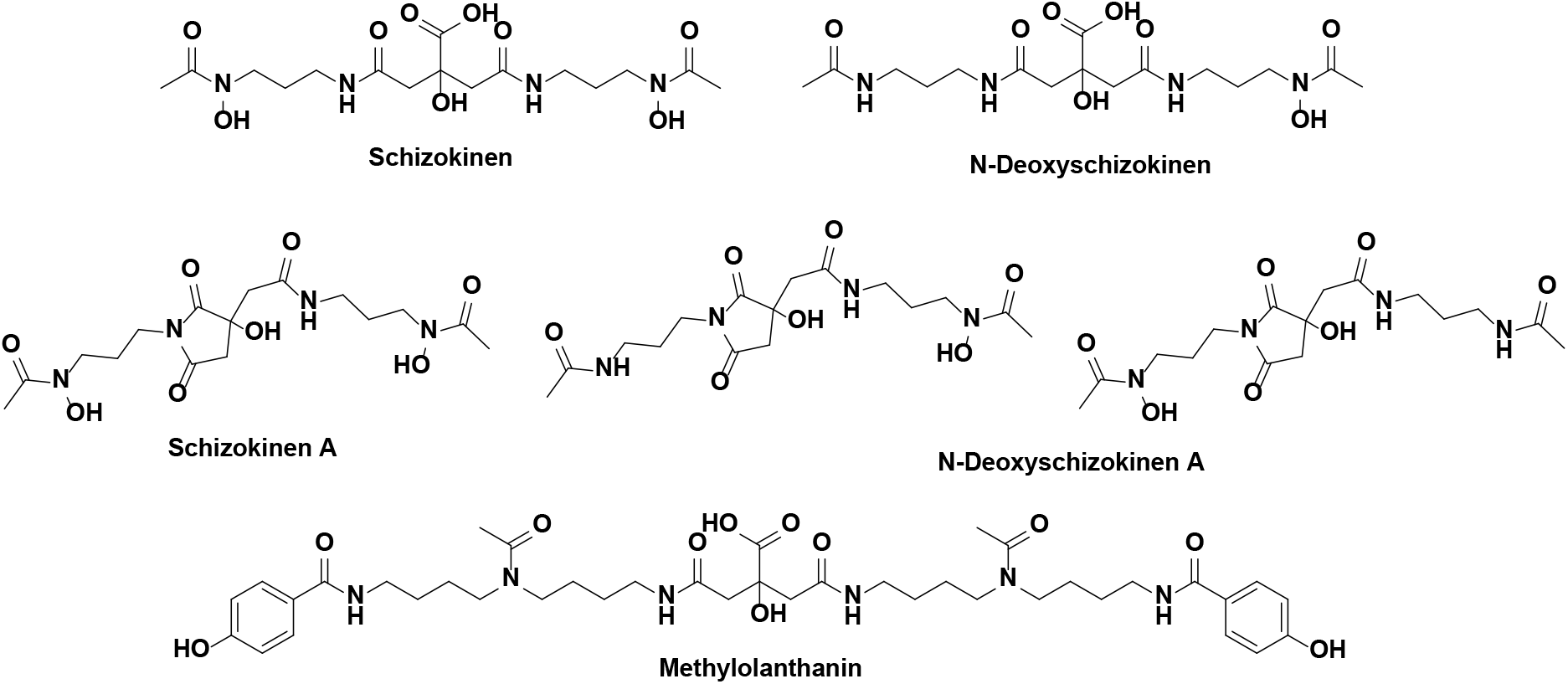
Schizokinen derivatives and methylolanthanin (MLL). For NDSA two variants are possible, but so far it has not been resolved which structure is the prevalent one, hence both are shown.

## Results and Discussion

The assignment that AM1 produces NDSA was based on another preprint of the same author reporting the development of a LC-MS method claiming to separate and detect four different schizokinen derivatives, at least one of which, was claimed to be found in AM1 supernatants.^[9]^ However, i) the molecule in question eluted *markedly* later than the other three schizokinen derivatives (approx. at 17 min versus 6-8 min in the chosen method), a shift far greater than the single-hydroxyl structural difference from schizokinen A suggests, ii) its measured accurate mass (found 387.1767) deviated from the value calculated for [C_16_H_26_N_4_O_7_+H]^+^ (387.1874) by −28 ppm, unreasonable for the HR-MS instrument reportedly used and well outside the deviations (between +3ppm and +5ppm) obtained for the other co-analyzed molecules within the standard: [Schizokinen +H]^+^ = [C_16_H_28_N_4_O_9_+H]^+^: found 421.1941, calc 421.1929, deviation 3ppm, [N-deoxyschizokinen +H]^+^= [C_16_H_28_N_4_O_8_+H]^+^: found 405.2002, calc 405.1980, deviation 5ppm, [Schizokinen-A +H]+ =[C_16_H_26_N_4_O_8_+H]^+^: found 403.1845, calc 403.1823, deviation 5ppm). Note that these three masses are consistently measured slightly (0.001-0.002 Da) *above* the calculations (due to small calibration imperfections in that mass range), while the measured peak at 387.1767 is significantly (>0.01 Da) *below* the calculation for [C_16_H_26_N_4_O_7_+H]^+^, i.e. the assigned sum formula is obviously wrong. iii) The MS/MS spectrum was dominated by a single fragment at *m/z* 105 whose accurate-mass formula is incompatible with the proposed structure and notably this fragmentation pattern is also very different to the other three schizokinen molecules in the standards. In addition, iv) the native, enzymatically made chelator is the open form (schizokinen, N-deoxyschizokinen), with the intact citrate core. A common observation in organic syntheses of citric acid-based siderophores is the formation of these imide side products, depending on the reaction conditions (e.g. choice of protecting groups and the pH).^[10–12]^ The imide A-form of schizokinen thus likely presents a downstream cyclization product in producing strains. Interestingly, the imide of the central citrate moiety in the siderophore staphyloferrin B has been shown to form under acidic conditions, but to isomerize to native staphyloferrin B under physiological conditions.^[13]^ So, a genuine schizokinen chelator producer should show the open form predominantly, with the imide as a minor derivative, certainly not the imide alone. Further, v) a fragment of *m/z* 77 was reportedly also found in the MS/MS spectrum at low intensity, textbook for an aromatic signature, of which N-deoxyschizokinen A is however devoid. The author even noted that the fragmentation pattern deviated from the other schizokinen species. In addition, as mentioned above a mass around *m/z* 105 was found. This could suggest the presence of a benzoyl ester moiety and points potentially towards a plastic contaminant.^[14]^ The used vials were not stated in the preprints, however we prepared LC-MS measurements in plastic vials (MN-702809) and glass vials (MN-702282) from Macherey-Nagel (MN) and a *m/z* 387.1802 feature can reproducibly be observed with the used plastic vials, but is absent when glass vials were used, identifying a leachable contaminant in all solvents tested (water, MeCN and MeOH). A similar introduced contaminant with a mass close to NDSA rather than a bacterially produced metabolite is thus likely (see Figure 2). This feature at *m/z* 387.1802 was found by manual EIC extraction of the putative mass of NDSA of *m/z* 387.1874 and applying a +/−30 ppm tolerance. We also detected this feature when we treated several polypropylene tubes in the lab with water or acetonitrile and measured the resulting solutions in glass vials, demonstrating the ubiquitous occurrence of this plastic-derived contaminant.

**Figure 2.**
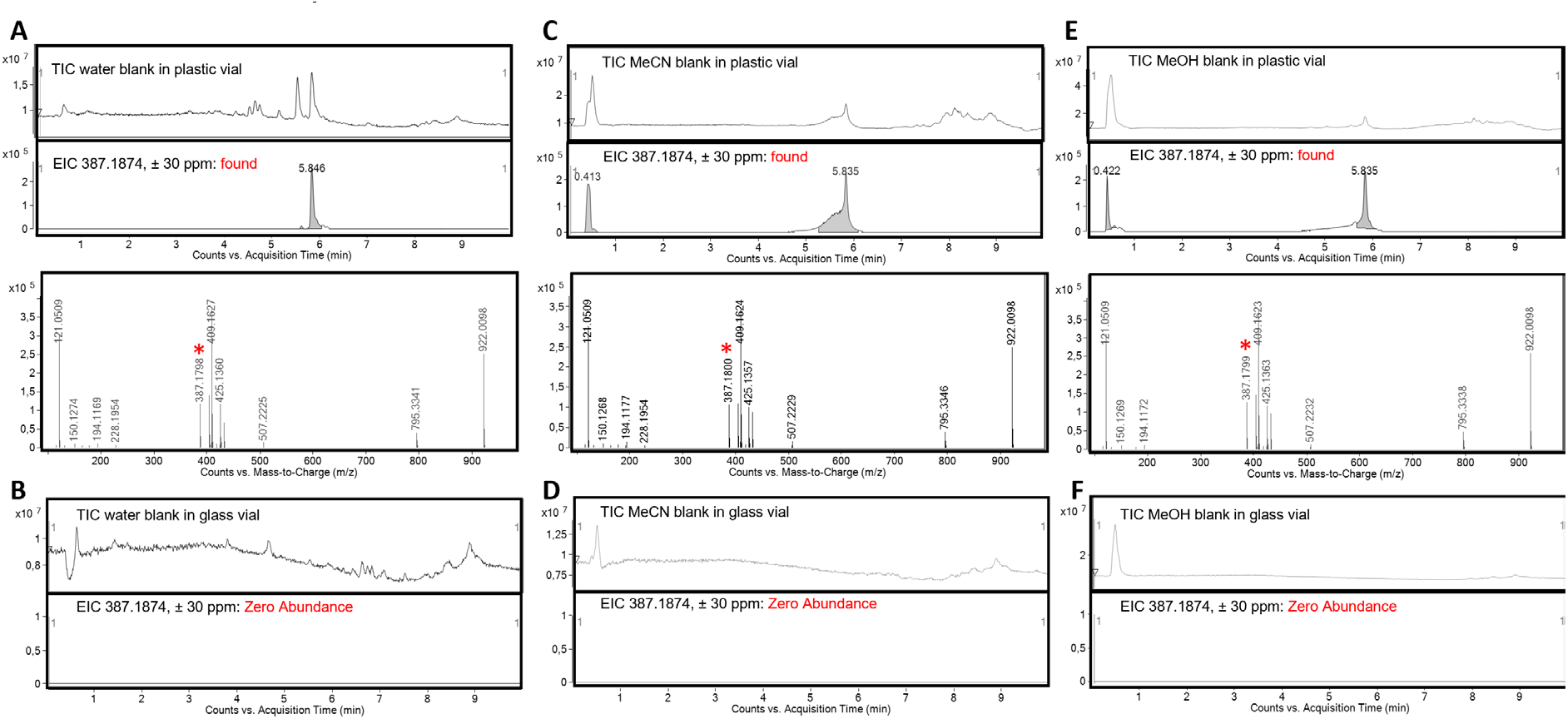
**A** Pure LCMS water in plastic vial vs. LCMS water in glass vial; injection volume: 50 µL. MN glass vial: No detection of *m/z* 387.1874. MN plastic vial: Detection of *m/z* 387.1874. **B** MeCN in plastic vial vs. MeCN in glass vial; injection volume: 50 µL. MN glass vial: No detection of *m/z* 387.1874. MN plastic vial: Detection of *m/z* 387.1874. **C** MeOH in plastic vial vs. MeOH in glass vial; injection volume: 50 µL. MN glass vial: No detection of *m/z* 387.1874. MN plastic vial: Detection of *m/z* 387.1874.

Next, we diluted schizokinen standard obtained by the same company as reported in the preprint with water to 10 µg/mL and measured the sample in a glass vial. With an injection volume of 1 µL the EIC of 387.1874 even with a tolerance of ± 30 ppm could not be detected (Figure 3B), Figure 3 shows this along with two schizokinen derivatives that were present in the standard mix. Schizokinen and schizokinen A could be identified via manual EIC *m/z* 421.1929 ± 5 ppm and *m/z* 403.1823 ± 5 ppm.

**Figure 3.**
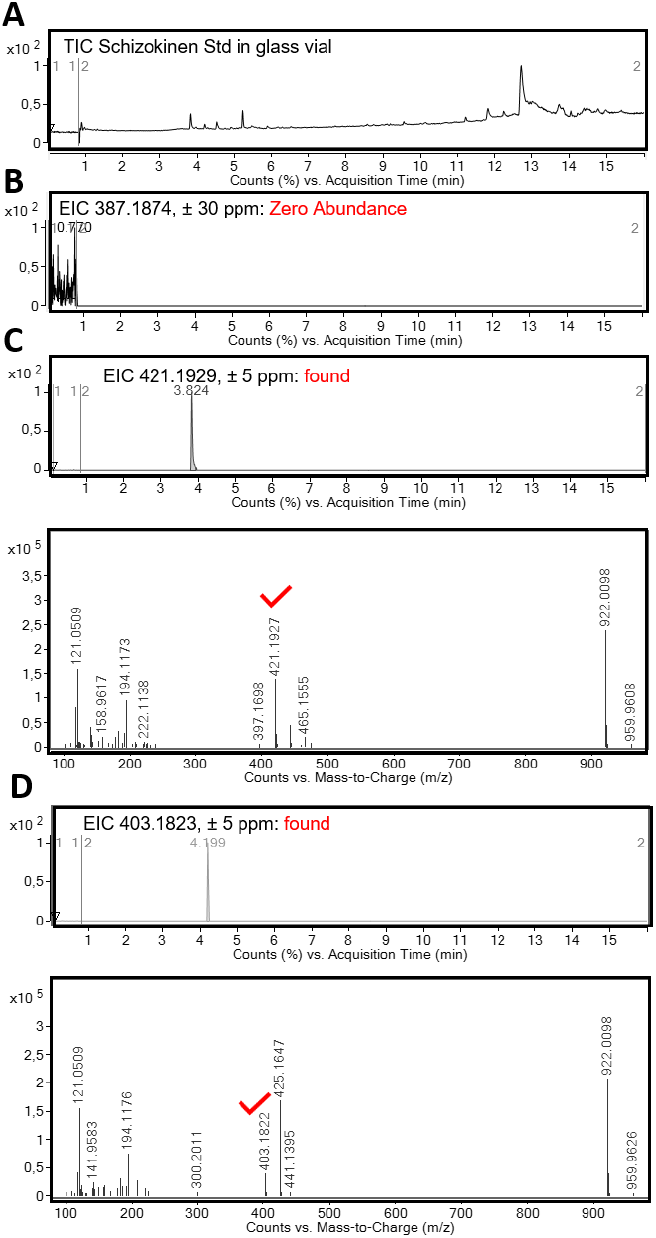
**A** TIC of Schizokinen standard measured in a MN glass vial. **B** N-deoxyschizokinen A (*m/z* 387.1874) could not be identified in the diluted standard measured in glass vials even with a threshold of ± 30 ppm. **C** Schizokinen and **D** schizokinen A could be identified via manual EIC 421.1929 ± 5 ppm and

And finally, vi) AM1 does to the best of our knowledge not have the necessary biosynthetic machinery to make schizokinen derivatives. Its citrate-methallophore cluster (*mll*) has been confirmed to make a petrobactin-type chelator with an aromatic residue (methylolanthanin).^[3,7]^ The claimed evidence for a second schizokinen derivative is thin, only a very low intensity mass spectrum without an accurate mass is presented, despite the capabilities of the instrument used in the study. Given the high concentration of the standard injected for the method development reported in the associated preprint^[9]^ (50 μL of a 0.1 mg/mL stock) this mass may have been derived from a contamination from the standard (memory effect) prior to measurements of the bacterial samples. Hence, the reported detection of schizokinen derivatives in *M. extorquens* AM1 should be regarded as a contamination derived artifact, and any conclusions that depend on its production (which have been done based on LC-MS analysis alone) should be revised accordingly. The positive arsenazo III assay reading in the study are most likely due to the chelator that AM1 actually produces: methylolanthanin.^[3]^ The plates for the CAS overlay assay reported in the paper were incubated for 30 days with the CAS layer, making any readout highly questionable. We therefore stress, there is no evidence that AM1 produces the siderophore N-deoxyschizokinen A.

## Conclusions

On the basis of the above discussion on the reported new chelator finding in strain AM1, most decisively the disappearance of a feature with similar mass in glass vials and the mass mismatch, the reported detection of *N*-deoxyschizokinen A in *M. extorquens* AM1 should be regarded as an artifact. Any conclusion that depends specifically on its production should be revised accordingly. More generally, we suggest that assignment of trace siderophore-like features in complex extracts be supported by glass-vial controls and solvent blanks, a reasonable accurate-mass threshold, and, where available, authentic standards and of course isolation or at least a second means of characterization.

## Methods and Materials

The same schizokinen standard as described in the preprint^[9]^ was used (obtained from BIOPHORE RESEARCH PRODUCTS, G. Winkelmann, Tübingen, Germany https://www.siderophore.com/) und a stock solution prepared in water (1 mg/mL) which was further diluted to 10 µg/mL for measurements. Plastic vials that were used were from Macherey-Nagel: 0.3 mL Polypropylene Snap Ring Vial N 11 outer diameter: 11.6 mm, outer height: 32 mm transparent, with inner cone, clear, MN-702809. Glass vials that were used were from Macherey-Nagel: Screw neck vial, N 9, 11.6x32.0 mm, 1.5 mL, flat bottom, clear, catalogue number 702282. Ultrapure water (type 1, pH 5.6, 18.2 MΩ·cm at 25 °C; *Merck Millipore®* Synergy® UV system) and LC-MS grade solvents (acetonitrile, methanol) and LC-MS grade formic acid (FA, *Sigma Aldrich*) were used if not stated otherwise.

**High-resolution mass spectrometry (HR-MS)** was measured on an *Agilent* 6530 C QTOF coupled to an *Agilent* 1260 Infinity II HPLC system (6530 C QTOF, G7115A DAD, G7116A, a G7167A, G7104C). Chromatograms and mass spectra were processed in MassHunter Qualitative Analysis software from *Agilent* version 10.0.

### Method for the plastic/glass vial comparisons

An *Agilent* Zorbax Extended-C18 (2.1 x 50 mm, 1.8 µm) column was used. Flow: 0.4 mL/min. Method: 0.5 min 95/5 water/acetonitrile, then in 6.5 min up to 5/95 water/acetonitrile, 3 min at 5/95 water/acetonitrile, followed by 2 minutes post-run. Source parameters (tuned in Extended Dynamic Range *m/z* 3200): positive mode; Gas Temp: 325 °C, Drying Gas: 8 L/min, Nebulizer 35 psi, Sheath Gas Temp 375 °C, Sheath Gas Flow 11 L/min; VCap 3500 V, Fragmentor 100 V, Skimmer 65 V, Oct 1 RF Vpp 750 V; mass range: *m/z* 100-1500; rate 2 spectra/s; time 500 ms/spectrum; transients/spectrum 4955; Reference masses used: 121.0509 and 922.0098; The column oven was set to 30 °C and the sample tray was cooled to 10 °C.

### Method for the schizokinen standard

An *Agilent* Poroshell 120 EC-C18 (3.0 x 150 mm, 2.7 µm) column was used. Used solvents: Flow: 0.7 mL/min. Method: 1 min 95/5 water/acetonitrile, then in 11 min up to 5/95 water/acetonitrile, held 4 min at 5/95 water/acetonitrile, followed by a 3 min post-run. Source parameters (tuned in Extended Dynamic Range *m/z* 3200): positive mode; Gas Temp: 250 °C, Drying Gas: 11 L/min, Nebulizer 45 psi, Sheath Gas Temp 350 °C, Sheath Gas Flow 12 L/min; VCap 3500 V, Fragmentor 100 V, Skimmer 65 V, Oct 1 RF Vpp 750 V; mass range: *m/z* 100-1700; rate 2 spectra/s; time 500 ms/spectrum; transients/spectrum 4950; Reference masses used: 121.0509 and 922.0098.

## Funding

The authors acknowledge financial support from the Deutsche Forschungsgemeinschaft (DFG) through the Collaborative Research Center “4f for Future” (CRC 1573, project number 471424360, A2) and European Research Council Starting grant Lanthanophore.

## Acknowledgements

The authors thank Harald Molitor from Macherey-Nagel for verifying the mass of 381.2 independently in their PP vials in their facility and Dr. Carl-Eric Wegner and Hannah Goerlach for helpful discussions.

## Notes

### Competing Interest Statement

The authors have declared no competing interest.

